# Designing a multi-epitope peptide-based vaccine against SARS-CoV-2

**DOI:** 10.1101/2020.04.15.040618

**Authors:** Abhishek Singh, Mukesh Thakur, Lalit Kumar Sharma, Kailash Chandra

**Author notes:** Corresponding authors: Dr. Mukesh Thakur, Scientist C - Centre for DNA Taxonomy & Coordinator - Centre for Forensic Sciences, Zoological Survey of India, New Alipore, Kolkata-700053, West Bengal, India, E. mail.

## Abstract

COVID-19 pandemic has resulted so far 14,395,16 confirmed cases with 85,711 deaths from the 212 countries, or territories. Due to multifacet issues and challenges in implementation of the safety & preventive measures, inconsistent coordination between societies-governments and most importanly lack of specific vaccine to SARS-CoV-2, the spread of Wuhan originated virus is still uprising after taking a heavy toll on human life. In the present study, we mapped several immunogenic epitopes (B-cell, T-cell, and IFN-gamma) over the entire structural proteins of SARS-CoV-2 and by applying various computational and immunoinformatics approaches, we designed a multi-epitope peptide based vaccine that predicted high immunogenic response in the largest proportion of world’s human population. To ensure high expression of the recombinant vaccine in *E. coli*, codon optimization and in-silico cloning were also carried out. The designed vaccine with high molecular affinity to TLR3 and TLR4, was found capable to initiate effective innate and adaptive immune response. The immune simulation also suggested uprising high levels of both B-cell and T-cell mediated immunity which on subsequent exposure cleared antigen from the system. The proposed vaccine found promising by yielding desired results and hence, should be tested by practical experimentations for its functioning and efficacy to neutralize SARS-CoV-2.

## Introduction

Coronavirus disease 2019 (COVID-19) is one of the most potent pandemics characterized by respiratory infections in Human. As per the latest report of World Health Organization (WHO) dated 10^th^ of April 2020, the worldwide coverage of COVID-19 exceeded 14,395,16 confirmed cases with mortality of 85,711 deaths from 212 countries, areas or territories [1]. The outbreak of COVID-19 emerged at Wuhan city of Hubei Province of P.R. China in the early December of 2019 [2], but the report of severe pneumonic patients with unknown etiology of symptoms was released by the Chinese Center for Disease Control on December 31^st,^ 2019 [3]. Later on, the novel Betacoronavirus (SARS-CoV-2) was confirmed as the causative agent for the clusters of pneumonic cases by the health professionals. Investigations on the origin of infections linked the SARS-CoV-2 virus to a seafood market in Wuhan city, China which got spread to a large extent of human population in China in a very short period of time. By the time, scientists/ health professionals paid a serious note to understand the mode of transmission of COVID-19, it had reached and infected several people in different countries with an insight into the escalated transmission rate. This led the WHO to declare COVID-19 as a Public Health Emergency of International Concern (PHEIC) on 30^th^ of January 2020 [4].

Until the outbreak of Severe Acute Respiratory Syndrome (SARS) in 2002 and 2003, coronaviruses were considered as a mild pathogenic virus causing respiratory and intestinal infections in animals and humans [5-9]. With a decade long pandemic of SARS, another potent pathogenic coronavirus, i.e., Middle East Respiratory Syndrome Coronavirus (MERS-CoV) appeared specifically in the Middle Eastern countries [10]. Extensive studies on these two coronaviruses have led to an understanding that they are genetically diverse but likely to be originated from bats [11, 12]. Apart from these two viruses, four other coronaviruses are commonly detected in Humans, viz., HKU1, 229E, OC43 and NL63 [13, 14]. Among all the coronaviruses reported so far, SARS-CoV, MERS-CoV, and SARS-CoV-2 have relatively high pathogenicity and escalated mortality rates. SARS CoV alone was responsible for 8422 positive cases and 919 probable fatalities over a vast population of 32 countries [15]. With the first mortality case reported in 2012, 2496 cases were found positive for MERS-CoV and 868 deaths were reported from 27 countries [15]. Both of these pandemics were considered as highly pathogenic in the last two decades until the outbreak of SARS-CoV-2 in December 2019. SARS-CoV-2 has already spread in 212 countries and caused 85,522 deaths in just last three and half months [1]. As in the current scenario, several countries are facing the most fatal problem of community spread of SARS-CoV-2 e.g. China, Italy, Spain, France, Iran, USA, India, Germany etc. Though, the estimated mortality rate due to SARS-CoV-2 is about 3.6% in China and about 1·5% outside China [16], the problem is grave and need to be respond urgently to curb the rapid loss of human life.

Coronaviruses are broadly divided into four genera, i.e., Alpha-CoV, Beta-CoV, Gamma-CoV and Delta-CoV, where Alpha-CoV and Beta-CoV are only reported to infect mammals but Gamma-CoV and Delta-CoV are also capable to infect both mammals and birds [17]. Extensive study of their biology suggests that they are non-segmented enveloped viruses with a positive-sense single-stranded RNA of a size range of 26-32 kb [18]. Some of the recent research suggested that SARS-CoV-2 has an identical genomic characterization as of Beta-CoV and thus the genomic structure of SARS-CoV-2 follows 5’-leader-UTR-replicase-S (Spike)–E (Envelope)-M(Membrane)-N (Nucleocapsid)-3’ UTRpoly (A) tail with the subsequent species-specific accessory gene at 3’ terminal of the viral genome [19]. The structural protein includes Spike/Surface protein (S), Envelop protein (E), Membrane protein (M) and Nucleocapsid protein (N). These Spike/Surface proteins are known to have crucial roles in survival and viral pathogenesis as the receptor binding capability and entry to the host cell is regulated by the influence of Surface protein [20]. The Envelop and Membrane proteins are responsible for viral assembly and Nucleocapsid protein is necessary for RNA genome synthesis [20]. This complex genetic makeup and moderate mutation rate of SARS-CoV-2 requires the strategic development of vaccine by targeting all the structural proteins. Unfortunately, there is no approved vaccine available as on date against SARS-CoV-2. However, a few studies have predicted peptide-based vaccine targets for specific glycoproteins [21, 22] but no study has covered all the structural proteins for vaccine designing. With the recent development of integrated bioinformatics and immunoinformatics approaches, vaccine designing and its specific application has become rapid and cost effective as with the computational simulations and predictions, the targeted immunogenic peptide can be designed and validated for its efficacy to work on human, even before having the vaccine in hand for practical exposure and experiments. However, conventional methods have simultaneously proven effective in vaccine designing with some limitation, e.g. sacrificing the whole organism or large protein residues which unnecessarily increases the antigenic load and probability of allergenicity [23]. We believe that an ideal multi-epitope vaccine candidate can enhance immune response and subsequently lower down the risk of re-infection by upraising the host immunogenicity. In this perspective, here we applied integrated approaches of vaccine designing by mapping B-cell, T-cell and IFN-gamma epitopes (humoral and adaptive immune response) of humans on the respective structural proteins based on several essential criteria.

### Methodology

Retrieval of SARS-CoV-2 protein sequences and antigenicity prediction All the protein sequences of SARS-CoV-2 available on NCBI (GenBank: MN908947.3) was retrieved and structural protein sequences were extracted for further analysis. We predicted antigenicity from each protein using the online server VaxiJen v2.0 [24] and proteins that showed antigenicity ≥ 0.4 in the virus category were subjected for further analysis.

### Prediction of physiochemical and secondary structural properties of the target proteins

The physicochemical properties of target proteins were examined using Expasy Protpram online server [25] and the conformational states, e.g. Helix, sheet, turn and coil were predicted using the online Secondary Structure Analysis Tool (SOPMA) [26] with default parameters.

### 3 Dimensional homology modeling and validation

Three-dimensional structure of target proteins was modeled using three homology modeling tools, i.e. I Tasser, Raptor-X and Phyre2 [27-29] and post processing to reduce any distortions in the modeled structure was done using the Galaxy refine server [30].The modeled structures were subjected to RAMPAGE server for Ramachandran plot analysis for quality check and validation[31 and visualized using Chimera 1.10.1 visualization system [32].

### Prediction of T-cell epitopes

#### Cytotoxic T-cell (CTL) epitopes

Nine residues long CTL epitopes recognized by HLA class-I supertypes, i.e., A1, A2, A3, A24, A26, B7, B8, B27, B39, B44, B58, B62 were predicted using NetCTL.1.2 server [33] considering the default values of weight on C terminal cleavage, weight on TAP transport efficiency and threshold for epitope identification. Further, another set of CTL epitopes recognized by HLA class I alleles like A*02:02, B*07:02, C*04:01, E*01:01, etc were identified by consensus method of Immune Epitope Database (IEDB) tool [34]. The identified epitopes were scrutinized based on the consensus score i.e. ≤2 and only those epitopes were considered as strong binders and selected for further analysis which were predicted by more than one allele of both the method used.

### Helper T-cell (HTL) epitopes

Fifteen residue long HTL epitopes recognized by HLA Class II DRB1 alleles were predicted using Net MHC II pan 3.2 server [35]. The thresholds for strong binder and weak binder were set as default. Further another set of HTL epitopes of 15 residues length recognized by HLA DR alleles were identified using the consensus method as implemented in the IEDB server [34] The epitopes with a percentile rank of ≤2 and predicted by more than one allele of both the methods were considered as a strong binder and scrutinized for further analysis.

### Identification of overlapping T-cell epitopes

Considering the fact that epitopes with affinity for multiple HLA alleles tend to induce relatively more immune response in the host cell, we scrutinized overlapping epitopes with affinity to both HLA class I and class II alleles for immunogenicity and allergen prediction using VaxiJen v2.0 server [24] and AllerTop v.2.0 [36] tool with an immunogenic threshold of 0.4. These epitopes would have high potential to activate both CTL and HTL cells.

### Identification of Continuous and Discontinuous B-cell epitopes

B-cell epitopes are short amino acid sequences recognized by the surface-bound receptors of B-lymphocyte. Their identification plays a major role in vaccine designing, and thus two types-Continuous and Discontinuous B-cell epitopes were predicted over the corresponding structural protein sequences. The continuous B-cell epitopes, present on the surface of proteins, were predicted using BCpred 2.0 [37] server and the discontinuous epitopes which were fairly large were predicted using Ellipro server [38]. The prediction parameters like Minimum score and Maximum distance (Anstrom) were set as 0.8 and 6, respectively.

### Identification of IFN-gamma epitopes

Since, IFN gamma is the signature cytokine of both the innate and adaptive immune system and epitopes with potency to induce IFN gamma could boost the immunogenic capacity of any vaccine. Thus, IFN gamma epitopes were predicted for each structural protein by the IFNepitope server [39].

### Characterization of predicted epitopes

#### Conservation analysis

Predicted T-cell and B-cell epitopes were submitted to the IEDB conservancy analysis tool [40] to identify the degree of conservancy in the structural protein sequences and the epitopes with 100% conservancy were selected for further analysis.

#### Population coverage and autoimmunity identification

Immunogenetic response of human population towards the selected overlapping CTL and HTL epitopes against their respective HLA genotype frequencies was predicted using IEDB population coverage analysis tool [41]. Those epitopes exhibiting 50% or more population coverage were scrutinized and to reduce the probability of autoimmunity, all the selected epitopes were subjected to BlastP search analysis against the Human proteome. Epitopes showing similarity to any human protein were excluded from the further analysis.

#### Interaction analysis of epitopes and their HLA alleles

The sequence of scrutinized epitopes was submitted to an online server PEPFOLD 3 [42] for 3D structure modeling. The structure of the most common HLA alleles i.e. HLA-DRB1 *01:01 (HLA class II) and HLA-A* 02:01 (HLA class I) in human population were retrieved from Protein data bank (PDB) with a PDB ID of 2g9h and 1QEW. Any ligand associated with the HLA allele’s structure was removed and energy-minimization was carried out. Thereafter, the modeled epitopes were docked with the corresponding HLA allele using a web server Patchdock [43], to study the interaction pattern of receptor and ligand. The parameters like clustering RMSD value and the complex type was kept as 0.4 and default. The result obtained from Patchdock was forwarded to the Firedock server [44] for the refinement of the best 10 models. Based on Global energy value, the complex with the lowest global energy was scrutinized and the corresponding CTL and HTL epitopes were selected for the final vaccine construct.

#### Construction and quality control of final multi-epitope vaccine

For the construction of a multi-epitope vaccine against SARS-CoV-2, we followed Chauhan et al. [23] with some modifications. The epitopes were finally selected based on meeting the following criteria: 1). epitopes must be immunogenic, non-allergic and overlapping with affinity to both HLA class I and class II alleles; 2).epitopes must be capable to activate both CTL and HTL cells and must have a minimum 50% of the population coverage; 3).epitopes should not be overlapping with any human gene and the predicted B-Cell epitopes should overlap with finalized CTL and HTL epitopes and should be present on the surface of the target protein. Based on these criteria, the HTL, CTL and IFN gamma epitopes were included in the final construct of the multi-epitope vaccine. The HTL and IFN gamma epitopes were linked by GPGPG linkers, whereas the CTL epitopes were linked by AAY linkers [23]. An adjuvant, 50S ribosomal protein L7/L12 (Locus RL7_MYCTU) with an NCBI accession no: P9WHE3 was added to the N terminal to enhance the immunogenicity of the constructed vaccine.

#### Antigenicity, allergenicity and physiochemical properties of vaccine

The antigenicity of the proposed vaccine was predicted using VaxiJen v2.0 server [24] a nd allergenicity was predicted using Allertop v.2.0 prediction tool [36]. The physicochemical property of the designed vaccine was predicted by submitting the final sequence of the vaccine to the Expasy Protpram online server [25].

#### Prediction of secondary structure

The secondary structure of the constructed vaccine was predicted using the SOPMA tool [26]. With the input of vaccine sequence and output width 70, the parameters like the number of conformational states, similarity threshold, and window width were set as 4 (Helix, Sheet, Turn, Coil), 8 and 17.

#### Modeling, refinement, and validation of vaccine construct

The final model of the constructed vaccine was prepared using a homology modeling tool SPARKS-X [45]. Among the top 10 predicted models, the model with the highest Z-score was selected and subjected for refinement using the Galaxy refine server [30]. Among all the refinement tools, Galaxy refine is one of the most efficient server according to CASP10 evaluation. Then after, we utilized ProSA-webtool [46] to assess the overall and local quality of the predicted model based on the Z-score. If the Z-score lies outside the range of native protein of similar size, a chance of error increases in the predicted structure. Further to check the overall quality of the refined structure of vaccine, Ramachandran plot analysis was carried out using the RAMPAGE server [31]. The final model of the vaccine construct was visualized using Chimera 1.10.1 visulaization system [32].

#### Prediction of Continuous and Discontinuous epitopes

Continuous and Discontinuous B-cell epitopes in the vaccine construct were predicted using BCpred 2.0 [37] and IEDB server [34], respectively. For effective designing of vaccine, the constructed vaccine should posses’ B-cell epitopes for potential enhancement of immunogenic reactions.

#### Molecular docking of constructed vaccine with TLR3 and TLR4 receptor

The 3D structures of human TLR3 and TLR4 were retrieved from protein data bank (PDB ID: 2A0Z and 2z63). Any ligand attached to the retrieved structures was removed and the receptor was subjected for model refinement using Galaxy refine server [30]. Molecular docking analysis was performed using the Cluspro v.2 protein-protein docking server [47] to analyze the interaction pattern of vaccine with TLR3 and TLR4. The server provides cluster scores based on rigid docking by sampling billions of conformation, pairwise RMSD energy minimization. Based on the lowest energy weight score and members, the final vaccine-TLR3 complex and TLR4 complex model was selected and visualized using Chimera 1.10.1 visulaization system [32].

#### *In silico* cloning and vaccine optimization

Efficiency of the multi-epitope vaccine construct in cloning and expression is of the utmost importance for vaccine designing. The final multi-epitope vaccine construct was submitted to the Java Codon Adaptation Tool (JCat) [48], where codon optimization was performed in *E. coli* strain K12. In addition to this, the parameters like Avoid rho-independent transcription terminators, Avoid prokaryotic ribosome binding sites, Avoid Cleavage Sites of Restriction Enzymes were selected for the final submission. The output of the JCat tool provided CAI value and GC content of the optimized sequence. Ideally, the CAI value should be greater than 0.8 or nearer to 1 [49], and the GC content should be in a range of 30 to 70% [50] to ensure that the vaccine has potential translation, stability and transcription efficiency. Further, an in silico cloning of a multi-epitope vaccine construct in *E. coli* pET– 28(+) vector was performed using the SnapGene 4.2 tool [51].

#### Immune simulations of vaccine construct

To characterize the real-life immunogenic profiles and immune response of the multi-epitope vaccine, C-ImmSim server [52] was utilized. Position-specific scoring matrix (PSSM) and machine learning are the two methods, based on which C-ImmSim predicts immune epitopes and immune interactions. It simultaneously simulates three different anatomical regions of mammals, i.e. Bone marrow, Thymus and tertiary lymphatic organs. To conduct the immune simulation, three injections at an interval of four weeks were administered with 1000 vaccine molecules per injection. The parameters like Random seed, simulation volume, and simulation step were kept as 12345, 10µl and 1050. Several studies suggested that the minimum interval between two injections should be kept four weeks and thus the time steps followed for three injections were 1, 84 and 168 where each time step is equal to 8 hours of real-life [53]. Since, several patients wordwide got recurrent infection of SARS-CoV-2, 12 consecutive injections were administered four weeks apart for the assessment of after effects of vaccine exposure on SARS-CoV-2.

## Results

### Target proteins and their antigenicity

We scrutinized target structural protein sequences, i.e. Surface, Membrane, Envelop and Nucleocapsid proteins from the whole-genome of SARS-CoV-2, 29903 bp ss-RNA (GenBank: MN908947.3). Antigenicity prediction from the structural protein sequences revealed that Envelop proteins has the highest antigenicity score 0.6025, followed by Membrane, Nucleocapsid and Surface protein (table 1).

**Table 1.** Details of target protein and their antigenicity score.

| **Structural Protein** | **Protein ID (NCBI)** | **Size (aa)** | **Antigenicity score** |
| --- | --- | --- | --- |
| Surface protein | QHD43416.1 | 1273 | 0.4646 |
| Membrane protein | QHD43419.1 | 222 | 0.5102 |
| Envelop protein | QHD43418.1 | 75 | 0.6025 |
| Nucleocapsid protein | QHD43423.2 | 419 | 0.5059 |

### Physiochemical and secondary structural properties

Theoretical estimates of isoelectric point (PI) suggested that all the structural proteins were basic in nature and their predicted instability indices were below 40 indicating their stable nature except the Nucleocapsid protein with an instability index of 55.09 (table S1). The predicted secondary structure characteristics of all the structural proteins showed varaible percentage of Alpha Helix, Extended Strand, Beta Turn and Random Coil (table S2).

### 3 Dimensional structure of target proteins

The refined finalized 3D structures obtained from Raptor-X was better than the models generated from I Tasser and Phyre2 as Raptor-X models showed least percentage of outliers and therefore selected for further analysis (figure 1). Details of various models of the structural proteins through Ramachandran plot analysis are given in table S3.

**Figure 1).**
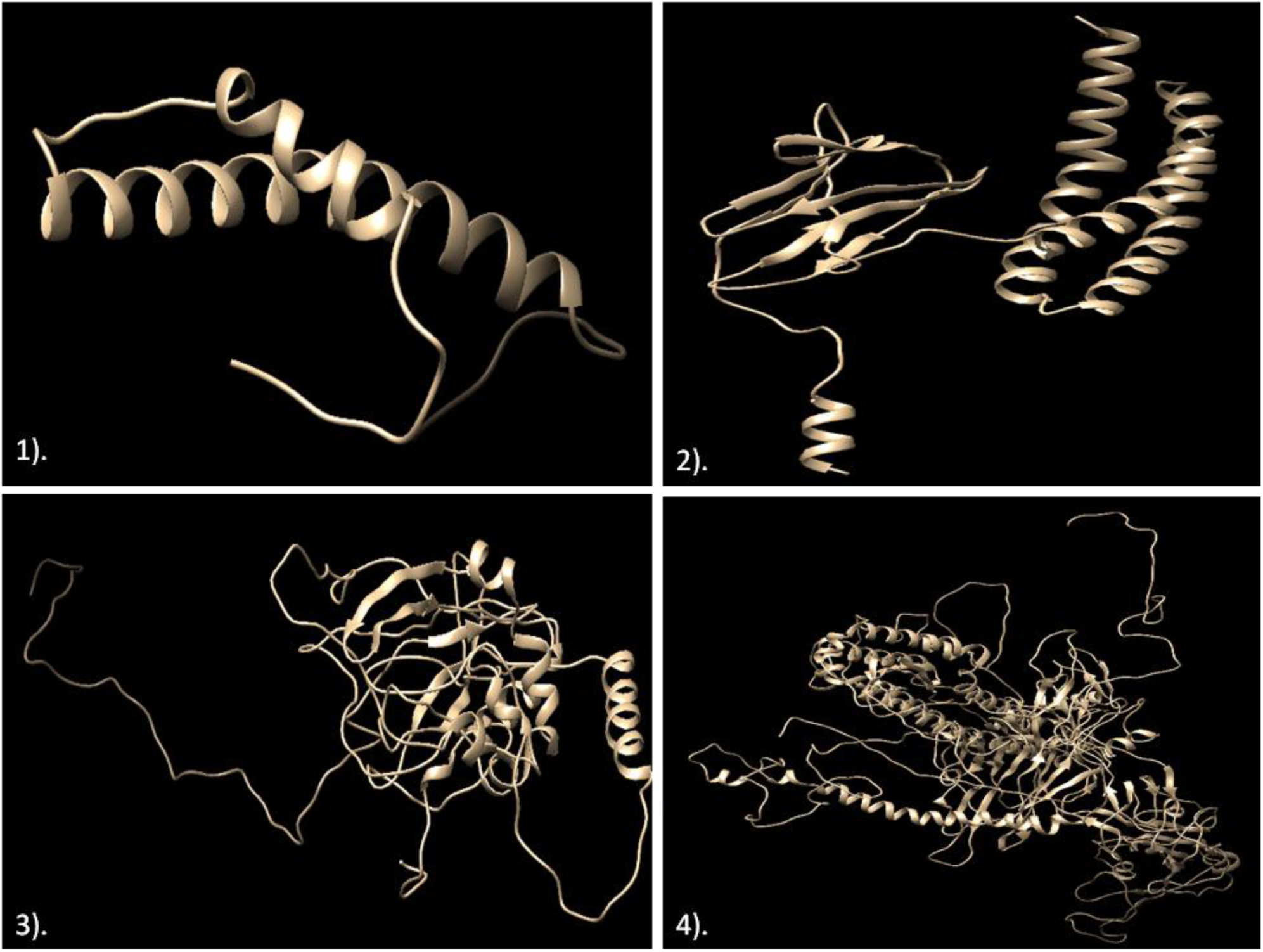
The 3D models of structural proteins of SARS- CoV-2.1). Envelop protein; 2). Membrane protein; 3). Nucleocapsid protein; 4). Surface protein

### Prediction of CTL and HTL epitopes

A total of 29 CTL epitopes in Envelop protein, 77 in Nucleocapsid, 340 in Surface and 89 epitopes in Membrane protein were predicted with strong binding affinity for multiple alleles. Similarly, 35 HTL epitopes in Envelop protein, 51 in Nucleocapsid, 252 in surface and 23 epitopes in Membrane protein were predicted. All these epitopes were strong binders and predicted percentile rank of ≤2.

### Identification of overlapping T cell epitopes

The CTL epitopes overlapping with HTL epitopes were scrutinized and subjected to immunogenicity and allergenicity prediction. In total, 16 CTL overlapped epitopes were predicted in Envelop protein, 8 in Nucleocapsid protein, 25 in Surface protein and 9 epitopes in Membrane protein (Table S4-S11). All these selected epitopes were predicted to be non-allergic with high antigenic scores.

### Identification of B-cell epitopes

The identified continuous B cell epitopes present on the surface of their respective proteins were selected based on their antigenic, non-allergic and overlapping nature in the finalized CTL and HTL epitopes. With these criteria, five epitopes were obtained in Surface protein, three in Nucleocapsid and one in Membrane protein (table S12). No epitope with such criteria was identified in Envelop protein. Further, one discontinuous epitope in Envelop protein, seven in Nucleocapsid, nine in Surface and four discontinuous epitopes were predicted in Membrane protein (table S13).

### Identification of IFN-gamma epitopes

The server IFNepitope predicted eight epitopes in Envelop protein, 39 in Membrane, 66 in Nucleocapsid and 281 in Surface protein. Based on antigenicity and non-allergenicity, eight epitopes were scrutinized in Envelop protein, 13 in Membrane, 43 in Nucleocapsid and 81 in Surface protein. To obtain the highest immunogenicity and reduce the overload of epitopes in vaccine construct, we selected one epitope from each protein with the highest antigenicity score and non-allergenicity (table S14).

### Characterization of predicted epitopes

#### Conservation analysis, Population coverage, and autoimmunity identification

The conservation prediction resulted that all the selected T cell and B cell epitopes were 100% conserved among the target structural proteins and population coverage prediction showed that five epitopes in Envelop protein, two in Membrane, two in Nucleocapsid and six epitopes in Surface protein covered more than 50% of worldwide population. No selected epitope with >50% population coverage showed homology with any human protein. Thus, altogether 30 epitopes, 15 each of CTL and HTL, met the above criteria were processed for the interaction analysis with the commonly occurring HLA alleles in the Human population (table S15).

### Interaction analysis of epitopes and their HLA alleles

Molecular interaction analysis of each selected CTL and HTL epitopes with their respective HLA alleles resulted in 10 predicted docked complex. Each complex showed different Global energy value. Out of the characterized 30 epitopes, the epitopes with highest negative value of Global energy (set threshold -40) were considered as effectively interacting epitopes with the corresponding HLA allele, and thus selected for final vaccine construct (table 2). This resulted in two CTL and HTL epitopes in Envelop protein, one in Membrane, two in Nuc leocapsid and two in Surface protein. For an ideal vaccine, the epitopes included should have potential affinity towards their respective alleles and thus greater the interaction between epitopes and their alleles, more is the affinity.

**Table 2.**
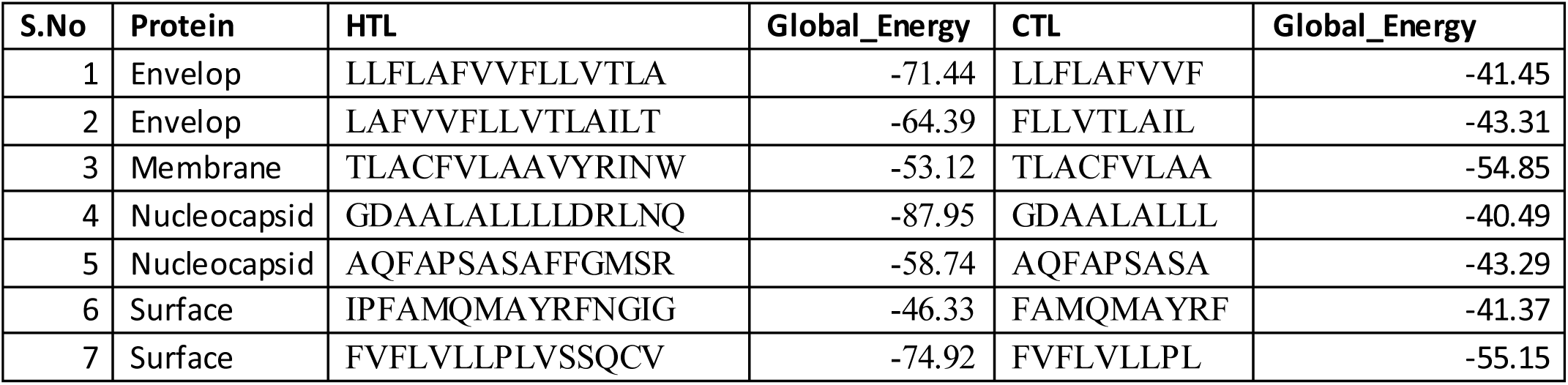
Final epitopes selected for multi-epitope vaccine construct after docking analysis with respective HLA alleles

### Construction of final multi-epitope vaccine

We selected seven CTL, seven HTL and four IFN gamma epitopes for construction of the multi-epitope vaccine (figure S1). The adjuvant was coupled by EAAAK linker with CTL epitope and subsequently, AAY linker was used to couple CTL epitopes and GPGPG linker was used to couple HTL and IFN gamma epitopes (figure 2). The final multi-epitope vaccine construct was composed of 430 amino acid residues which was then validated for antigenic, allergenic and physiochemical properties.

**Figure 2.**
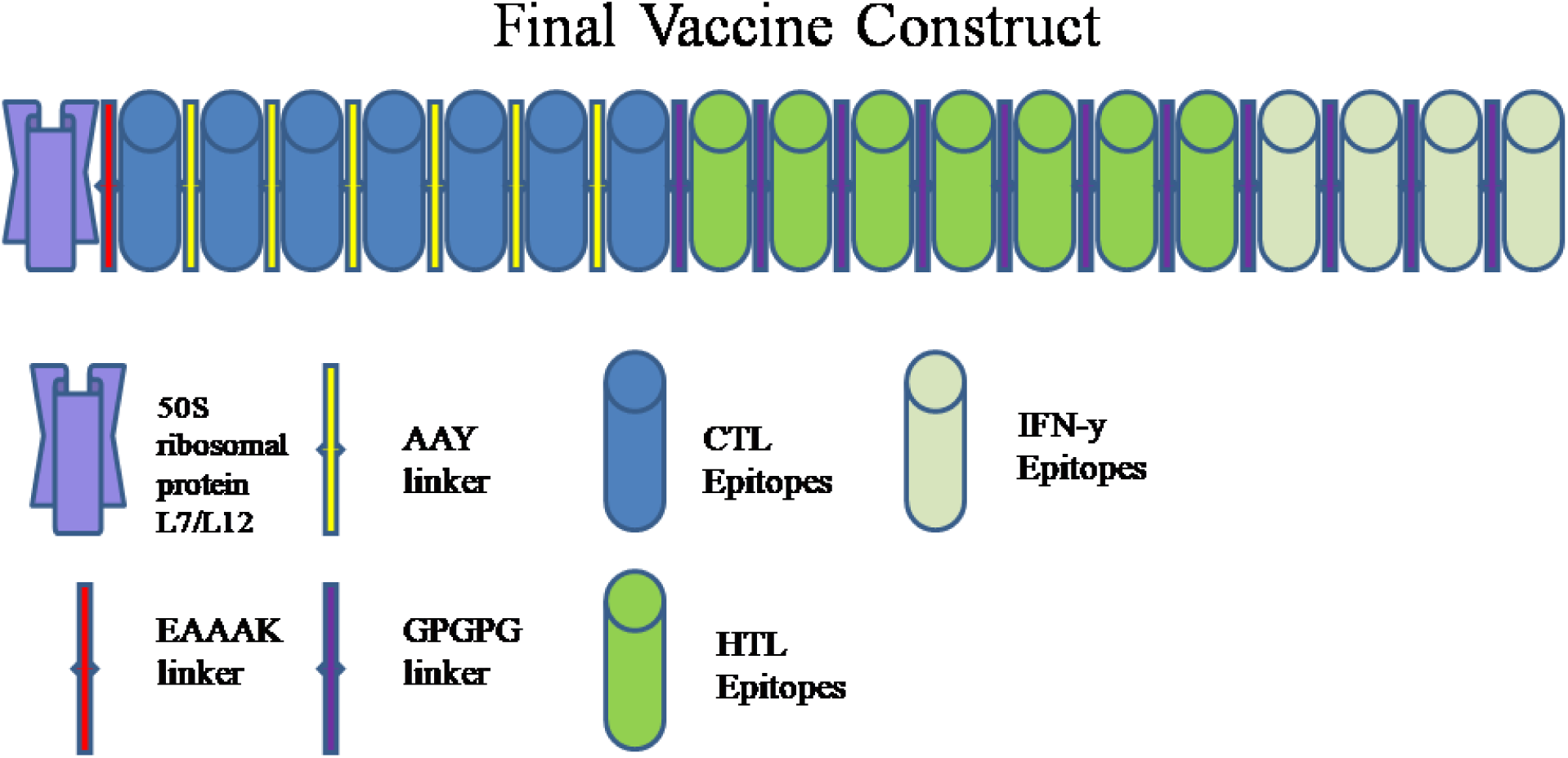
Schematic diagram of final vaccine construct

### Evaluation of antigenicity, allergenicity, physiochemical properties and secondary structure of the vaccine construct

The multi-epitope vaccine construct found to be immunogenic (Ag score-0.627), non-allergic with 45.131 KDa predicted molecular weight. The prediction suggested that vaccine construct was stable and basic in nature with instability index 23.46 and theoretical PI 9.01. The half-life of the vaccine in mammalian reticulocytes (in vitro) was 30 hrs, while it was about 20 hrs in yeast (in-vivo) and 10 hrs in *E. coli* (in vivo).

The predicted secondary structure of the vaccine constructed consisted 42.33% alpha-helix, 22.09% extended strand, 4.42% beta-turn and 31.16% random coil.

### Modeling, refinement, and validation of vaccine construct

Of the top 10 predicted models, the model with the Z-score of 7.61 with template 1dd3A was selected for refinement. The ProSA-web tool assessed the overall and local quality of the refined model with a Z-score of -0.47 which was close to the range of native protein of similar size (figure 3). Further, the Ramachandran plot analysis revealed 92% of residues in the favored region, 6.1% in the allowed region and 1.9% outliers. This signified that the quality of the predicted vaccine construct was adequate for further analysis.

**Figure 3.**
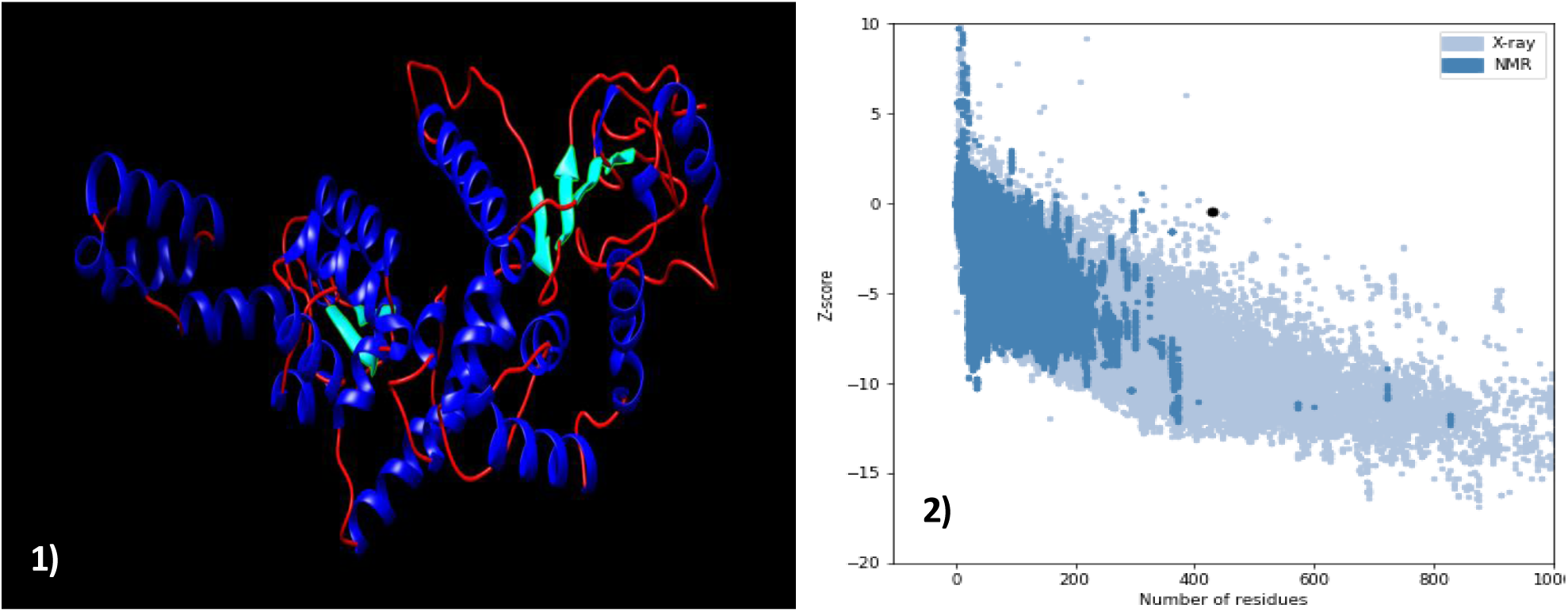
1). 3 dimensional structure of final vaccine construct, 2). ProSA validation of predicted structure with Z score of -0.47.

### Prediction of Continuous and Discontinuous epitopes

BCpred 2.0 server predicted four continuous epitopes i.e. AKILKEKYGLD, EILDKSKEKTSFD, LKESKDLV and VPKHLKKGLSKEEAESLKKQLEEV on the surface of predicted vaccine (figure S2), while IEDB predicted six discontinuous epitopes (figure S3) with a score higher than 0.8 (Table S16).

### Molecular docking of constructed vaccine with TLR3 and TLR4 receptor

Cluspro v.2 predicted 30 models each of vaccine receptor TLR3 complex and TLR4 complex with their corresponding cluster scores (Table S17-S18). Among these models, the model number 2 in TLR3 complex and model number 1 in TLR4 complex were selected as a best-docked complex with the lowest energy score of -1199.1 with 46 members (TLR3) and lowest energy score of -1229.9 with 79 members (TLR4). This signifies potential molecular interaction between predicted vaccine construct with TLR3 and TLR 4 receptors (figure 4).

**Figure 4.**
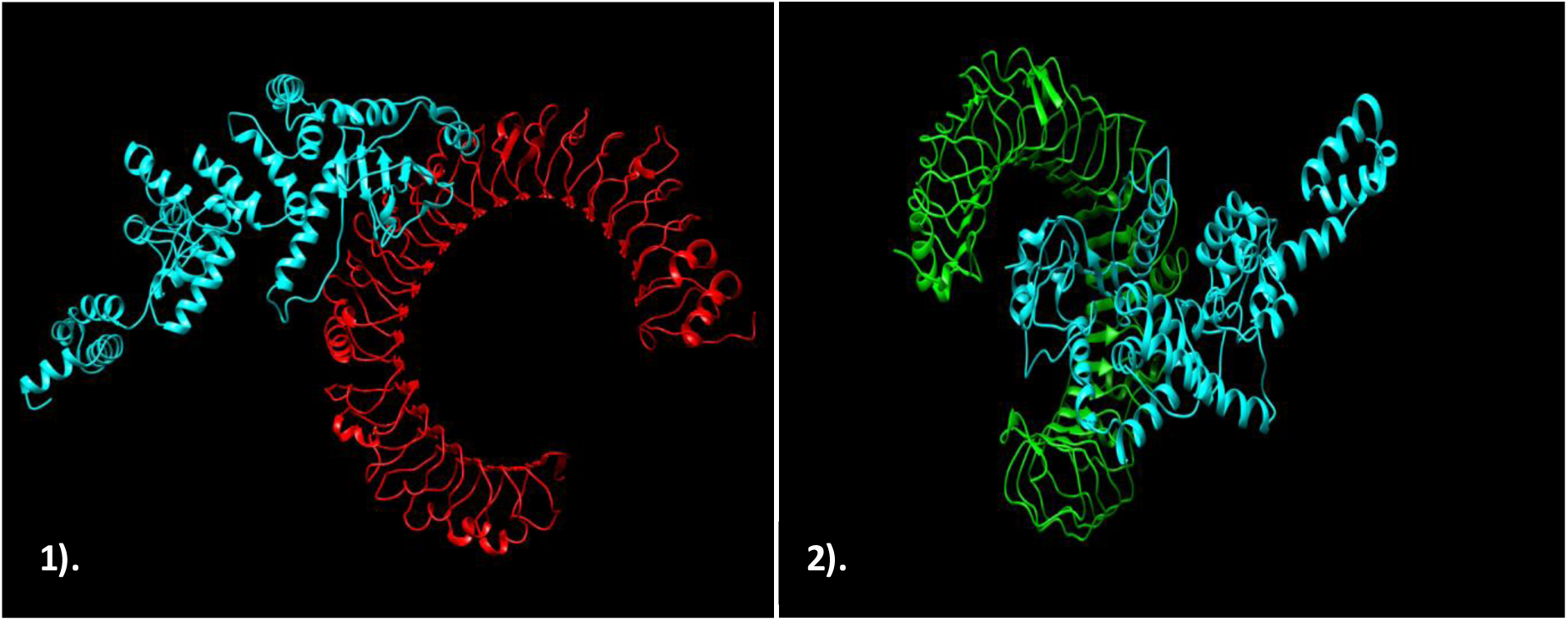
The interaction pattern of designed vaccine with TLR3 and TLR4. **1)**. Vaccine (Cyan) docked with receptor TLR3 (Red). **2)**. Vaccine (Cyan) docked with receptor TLR4 (Green).

### *In silico* cloning and vaccine optimization

The Java Codon Adaptation Tool optimized the codon usage of the vaccine and produced an optimized codon sequence of length 1290 nucleotides. The CAI value of optimized sequences was 0.95, close to 1 and the GC content was found to be 54.41 %, fall between the range of 30 to 70%. Therefore, the adaptation was satisfactory indicating potential expression of vaccine construct in host *E*.*coli*. Later on, the restriction sequence of Xho I and Not I restriction enzymes were added to N and C terminal of adapted codon sequence. Further, the adapted sequence was also cloned in pET– 28(+) vector using SnapGene tool (figure 5).

**Figure 5.**
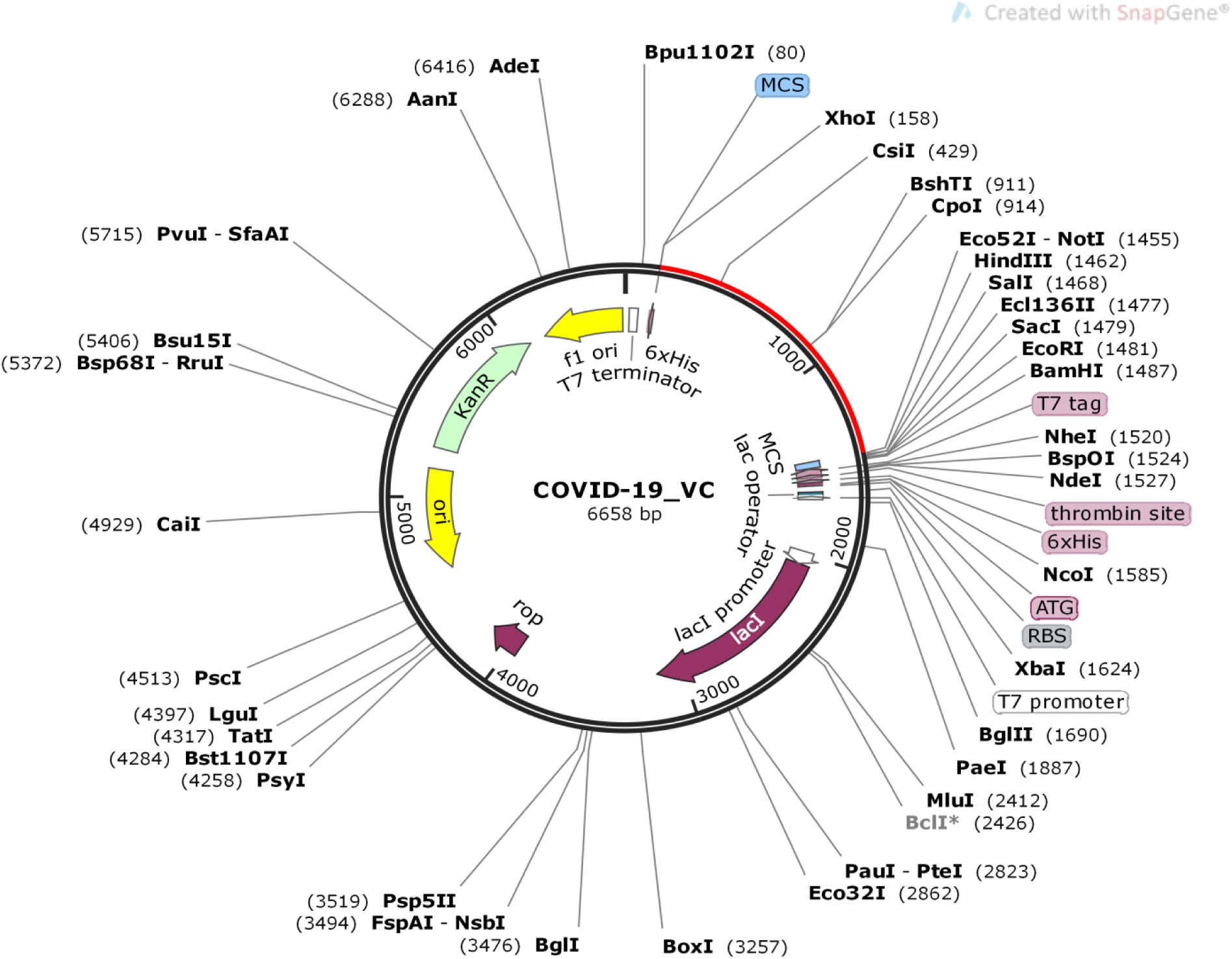
In silico cloning of vaccine. The segment represented in red is the multi-epitope vaccine insert in pET– 28(+) expression vector.

### Immune simulations of vaccine construct

C-ImmSim server-generated immune response was consistent with the actual immune response as depicted by the results (figure 6 and S4). The primary response was generated as marked by the increase in the level of IgM. Similarly, the secondary response was characterized by an increased level of IgM+IgG, IgG1+IgG2, IgG1, IgG2, and B-cell populations. On the subsequent exposure of vaccine, a decrease in the level of antigens was observed indicating the development of immunogenic response in the form of immune memory. Both the T cell populations i.e. CTL and HTL developed increased response corresponding to the memory cells signifying the immunogenicity of T cell epitopes included in the vaccine construct. Increased activity of macrophages was also observed at each exposure, whereas the activity of NK cells was observed consistent throughout the period. A significant increase in the level of IFN gamma, IL-10, IL-23, and IL-12 was also observed at subsequent exposure (figure 6). After the repeated exposure of vaccine construct through 12 injections after a regular interval, a remarkable increase in the level of IgM+IgG, IgG1+IgG2. Though, IgG1 increased slightly but IgM and IgG2 increased consistently. A surge in the memory corresponding to B cell and T cells was observed throughout the exposure while the level of IFN gamma was consistently high from first to last exposure (figure S4). This signifies that the vaccine proposed in this study has generated a strong immune response in case of short exposure and immunity increases even on subsequent repeated exposure.

**Figure. 6.**
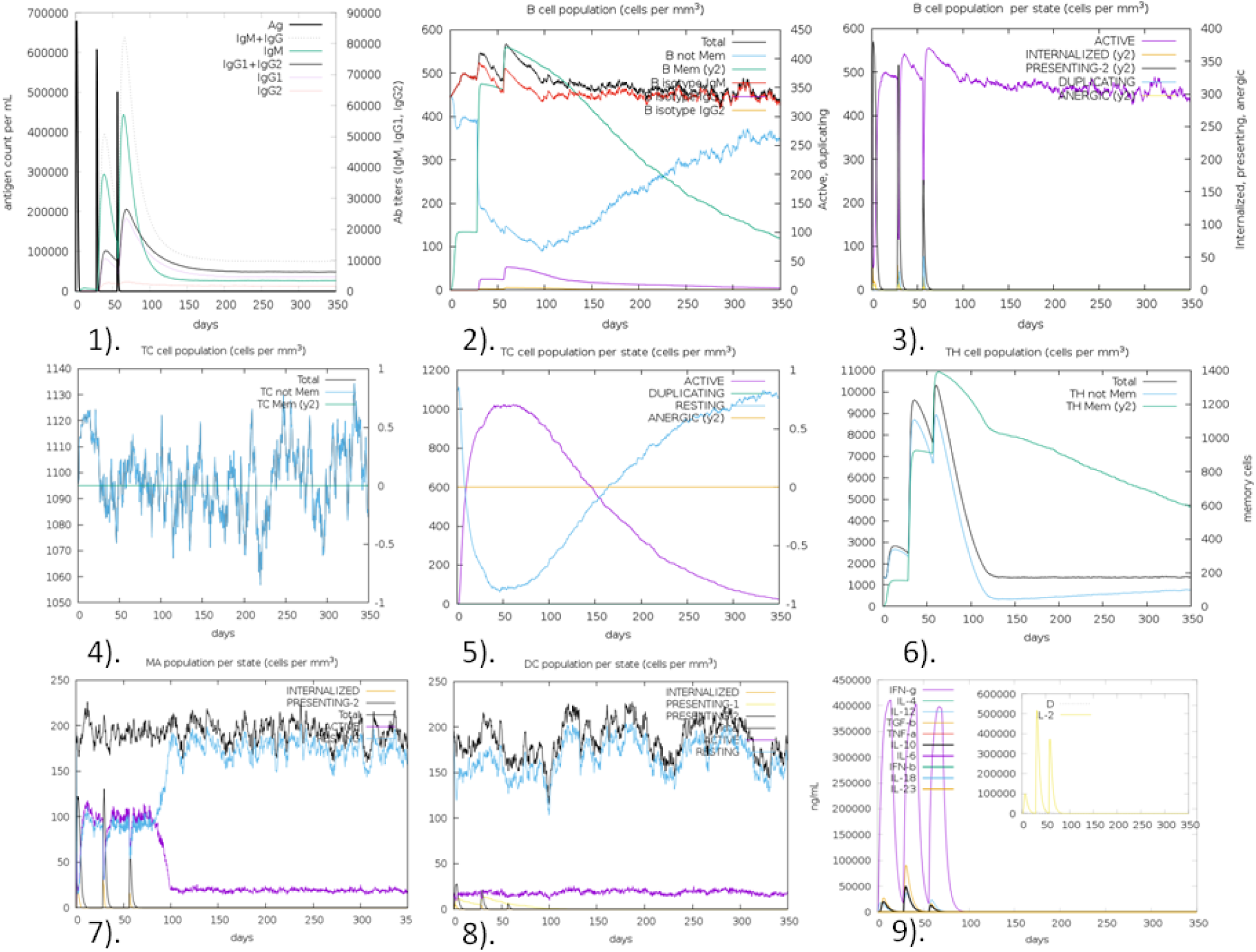
In silico simulation of immune response using vaccine as an antigen after subsequent three injections. 1). Antigen and Immunoglobins. 2). B-cell population. 3). B-cell population per state. 4). Cytotoxic T-cell population. 5). Cytotoxic T-cell population per state. 6). Helper T-cell population. 7). Macrophages population per state. 8). Dendritic cell population per state. 9). Cytokine production.

## Discussion

The recent outbreak of SARS-CoV-2 in Wuhan city of China raised several questions regarding the susceptibility of Humans against these novel pathogens. As predicted earlier about the re-emergence of novel variants of SARS-CoV [54], SARS-CoV-2 proved to be more lethal as compared to the previous variants in terms of wide spread of virus in a short period of time. This escalated rate of transmission initiated a pursuit for a vaccine development against SARS-CoV-2, which is currently a worldwide pandemic with over 1.43 million cases and 85, 711 deaths [1]. With strict and comprehensive measures, vaccine development and its application could play a key role in eliminating the virus from the Human population or to restrain the spread among individuals and different populations. Thus several efforts are being made to address the challenge that appeared in the current scenario with appreciable advancements in the understanding of virus biology and its etiology. The lack of knowledge regarding the response of the immune system against viral infection is one of the major limitations in the path of vaccine development for SARS-CoV-2.

This study is the piolt attempt in describing the potential immunogenic target over the structural proteins and proposes a novel multi-epitope vaccine construct, by providing new rays of hope in the initial phase of vaccine development. Certain criteria like poor antigenicity, allergenicity, low affinity towards the immune cells, autoimmunity and oversize that could influence the effectiveness, have been evaluated on the proposed vaccine construct following various computational and immunoinformatics approach. The retrieved structural proteins and their antigenicity score suggested that the Envelop protein is the most potent protein to generate immune response considering its role in viral assembly. Since, immunity against any pathogen is prominently dependent, how it gets recognized by B cells and T cells. We identified epitopes corresponding to B cells and T cells in each structural protein so that both humoral and cellular immunity can be induced with the exposure of vaccine construct. The proposed vaccine construct is designed based on the epitopes that have been selected with the most robust criteria,e.g. their nature of antigenic, non-allergic, 100% conserved among the target proteins, their affinity for multiple alleles, no homology with any of the human proteins, their worldwide coverage of human population (>50% of worldwide population covered) and effective molecular interaction with their respective HLA alleles. Further,, IFN gamma epitopes which are equally effective in immune regulatory, antiviral and antitumor activities, the final vaccine construct was designed with also due considering of the IFN gamma epitopes to increase the immune recognition ability. The designed vaccine of 430 amino acid residues has a molecular weight of 45.131K Da, also falls in the defined range of average molecular weight of a mullti-epitope vaccine. The instability index and theoretical PI also suggested that the vaccine is stable and basic in nature and the estimated half-life suggested that the recognized peptide does not possess a short half-life and would remain viable for a span adequate to generate a potential immune response. Vaccine model refinement and validation indicated that quality of the predicted model was good as more than 90% residues were in the favored region [55]. The B-cell continuous epitopes predicted over the surface of the vaccine suggested that the vaccine is capable to get recognized by B-cells indicating its effectiveness. TLR 3 and TLR 4 have proven recognition capability in both SARS-CoV and MERSCoV. Considering the similar genome characterization of SARS-CoV and SARS-CoV-2, the molecular interaction of vaccine with TLR 3 and TLR 4 through docking analysis suggested that the constructed vaccine has a significant affinity towards the toll-like receptors to act as sensor for recognizing molecular patterns of pathogen and initiating immune response. Thus the vaccine TLR complex is capable of generating an effective innate immune response against SARS-CoV-2. Further, Codon adaptation improved the expression of the recombinant vaccine in *E coli*. Strain K12 with significant codon adaptation index and GC content indicating elevated expression level. After successful cloning of the recombinant vaccine in the pET– 28(+) vector, the simulation based generated immune response suggested that the primary and secondary immune response will be coherent with the actual expected response. Subsequent exposure for three injections indicated the potential capability of vaccine for generating memory cells and a higher probability of significant immune response on later exposure to coronavirus. The consistent high level of IFN gamma also supported the activation of humoral immunity. To understand the immunogenic potential of vaccine, repeated exposure in the form of 12 injections was administered and the result suggested the consistent generation of a strong immune response. With all these immunoinformatics approaches, the vaccine designed against SARS-CoV-2 showed promising results in inducing immune response.

Without proper control methods or vaccines, it is difficult to curb coronavirus pandemic given the current situation. However, several medications have been tested against SARS-CoV-2, but none of them showed complete effectiveness. In this study, a vaccinomics approach was carried out to design a multi-epitope vaccine against SARS-CoV-2 using several in silico and immunoinformatic approaches. Based on the computational and immunoinformatics approaches, we propose that the designed vaccine has all the potential to induce both the innate and adaptive immune systems and can neutralize the SARS-CoV-2, and therefore, an experimental validation of the designed vaccine must be undertaken by the professional for reaching a conclusion for its safety, efficacy and success.

## Acknowledgements

We thank Zoological Survey of India, Kolkata for providing access to the computational support, working space and facilities to undertake the present study.

## Notes

### Competing Interest Statement

The authors have declared no competing interest.

